# ^13^C-Metabolic Flux Analysis of Substrate Co-Metabolism in the Methylotrophic Acetogen *Eubacterium limosum*

**DOI:** 10.64898/2026.10.12.758492

**Authors:** Kathryn O. Hoyt, William Gasparrini, Justin A. Hayes, Benjamin M. Woolston

## Abstract

Acetogens are promising microbial catalysts for the sustainable production of fuels and chemicals, metabolizing a range of renewable single-carbon (C1) compounds with high energetic efficiency. The major challenge is their energy-limited anaerobic lifestyle, which results in slow growth and a limited product spectrum on individual C1 substrates. We previously showed in the acetogen *Eubacterium limosum* that co-metabolism of glucose-methanol or CO-methanol mixtures results in substantial improvements in growth and acetate productivity. To investigate the underlying mechanism(s), here we conducted ^13^C metabolic flux analysis (^13^C-MFA) of *E. limosum* grown on a glucose-methanol mixture, as well as in single-substrate controls. The results suggest that the main benefit of glucose cofeeding is not provision of additional ATP, but rather a reduction of flux through the ferredoxin-dependent and kinetically challenging reactions of the Wood-Ljungdahl pathway. Subsequent RNAseq analysis showed no significant differences in the expression level of genes involved in methanol metabolism between cofeeding and methanol-only controls, suggesting this reduction of flux is controlled post-transcriptionally. Most surprisingly, we found that under all growth conditions, flux through pyruvate:ferredoxin oxidoreductase (PFOR) was in the oxidative direction, with cysteine (added as a reducing agent) providing all required gluconeogenic carbon flux under CO_2_-fixing conditions. This work provides critical insights into the metabolic network of an emerging model acetogen that will facilitate future metabolic engineering efforts to channel flux to value-added products, and highlights the under-appreciated role of cysteine as a carbon source in acetogenic fermentations.

**Highlights:**

- ^13^C MFA was conducted for *E. limosum* grown on methanol, glucose, or cofed.
- Cultures reached metabolic and isotopic stationarity in the AneVO chemostat.
- Flux distributions showed significant contributions of carbon from cysteine reducing agent.
- Cofactor analysis reveals cofeeding benefit is not simply additional ATP

## 1. Introduction

Single-carbon (C1) compounds have emerged as promising substrates for renewable biofuels and biochemical production, offering a route to displace petrochemical production while valorizing waste carbon resources (Aggarwal et al., 2023; Bachleitner et al., 2023; Jones et al., 2024; Lee et al., 2019). C1 feedstocks are abundant and do not compete with food systems, since they derive from industrial waste gases, lignocellulosic biomass (Daniell et al., 2012), and the electro- or thermochemical reduction of CO_2_, rather than from food crops (Dürre and Eikmanns, 2015). Gaseous C1 streams containing CO₂/H₂ or CO can be fed directly to microbial catalysts (Liew et al., 2022, 2016). Alternatively, CO_2_ can be converted to the more reduced, easier-to-store liquids methanol and formic acid (Jouny et al., 2018), each of which is itself a viable substrate for microbial upgrading (Cotton et al., 2020).

Acetogenic bacteria are attractive biocatalysts for C1 conversion (Bae et al., 2025, 2024; Da Silva Lameira et al., 2025; Gasparrini et al., 2025; Jin et al., 2020). Their appeal rests on the unusually high efficiency of their metabolism. Growth proceeds anaerobically through the Wood–Ljungdahl pathway (WLP, or reductive acetyl-CoA pathway), which channels the bulk of assimilated carbon into secreted products like acetate and butyrate, yielding carbon and energy conversion efficiencies approaching 90%, (Claassens et al., 2019; Fast and Papoutsakis, 2012). Among microbial methanol utilizers, methylotrophic acetogens are distinctive: they are the only known methylotrophs whose pathway results in net CO₂ consumption (Kremp et al., 2018; Sanford and Woolston, 2022a). *Eubacterium limosum* is one such organism and, unlike many acetogens, is genetically tractable (Bae et al., 2026; Flaiz et al., 2021; Jin et al., 2025; Sanford et al., 2024, 2023; Sanford and Woolston, 2024, 2022b; Shin et al., 2023, 2019), making it an attractive model for dissecting C1 metabolism and a promising chassis for engineered production.

The anaerobic lifestyle of acetogens comes at a cost. Across C1 substrates, the WLP generates less than one mole of ATP per mol of acetate produced (Schuchmann and Müller, 2014). This energy limitation results in long doubling times, low cell densities, and a constrained product spectrum (Lee et al., 2022; Poehlein et al., 2024). One strategy to relieve this energy limitation is substrate cofeeding, in which a second carbon source is supplied alongside the C1 substrate to accelerate growth or broaden the product spectrum (Liu et al., 2020). In *E. limosum* specifically, early work reported a 7.5-fold increase in specific growth rate and a transient 33% increase in specific methanol uptake rate when glucose was co-fed with methanol, relative to methanol alone (Loubiere et al., 1992). We recently validated this growth benefit in fed-batch culture, observing increases in growth rate, cell density, and productivity during glucose–methanol and CO–methanol co-metabolism (Hoyt et al., 2026).

Despite the apparent generality of the cofeeding benefit across acetogens, the mechanism(s) responsible for it remains unresolved. The prevailing hypothesis attributes the cofeeding benefit to supplemental ATP generated from the sugar co-substrate by glycolysis, which relieves the energetic bottleneck of C1 growth (Fast et al., 2015; Lee et al., 2022; Liu et al., 2020; Neuendorf et al., 2021). An alternative, less-explored possibility is that the second substrate redistributes carbon flux so as to reduce demand on kinetically or thermodynamically difficult reactions—for example, supplying biomass precursors through glycolysis rather than gluconeogenesis (Le Bloas et al., 1993; Loubiere et al., 1992). Chief among these difficult reactions is pyruvate:ferredoxin oxidoreductase (PFOR), which acetogens must run in the thermodynamically and kinetically unfavorable reductive (carboxylating) direction to generate pyruvate from acetyl-CoA and CO₂ during C1 growth (Furdui and Ragsdale, 2000). Correspondingly, PFOR is highly expressed in *E. limosum* (Song et al., 2018). Because cells adjust metabolite and enzyme levels to keep pathway free energies negative (Noor et al., 2014), reducing, reversing or eliminating flux through such reactions could substantially ease the burden of C1 growth. Distinguishing between the energetic and flux-redistribution hypotheses requires direct, quantitative measurement of intracellular carbon flux.

^13^C metabolic flux analysis (^13^C MFA) is the gold standard for quantification of in vivo carbon fluxes (Long and Antoniewicz, 2019), but applying it to C1 metabolism poses a challenge: growth on a single C1 substrate produces a uniformly labeled metabolome that carries little positional isotopic information. Cofeeding two distinctly labeled substrates overcomes this limitation (Hoyt and Woolston, 2022), making co-metabolism not only a growth strategy, but also a feature that renders acetogenic flux analysis tractable. Indeed, ¹³C-MFA has been applied to the acetogen *Clostridium ljungdahlii* during mixotrophic growth on glucose, CO_2_, and H_2_ (Dahle et al., 2022), providing useful frameworks for modeling flux in related Clostridia. A further, often-overlooked consideration in acetogen metabolism is the role of cysteine. Defined acetogen media frequently include cysteine as a reducing agent (Kountz et al., 2020; Mariotto et al., 1989), and its potential role as a carbon- or sulfur source is typically assumed to be negligible. However, cysteine metabolism has recently been demonstrated in *Moorella thermoacetica* (Göbbels et al., 2021) and *E. limosum* (Cao et al., 2026), warranting explicit accounting in flux analysis.

Here we performed steady-state ^13^C MFA for *E. limosum* grown on methanol/CO_2_, glucose/CO_2_, and methanol /CO_2_ cofed with glucose to resolve the carbon flux distributions under each condition and test the competing hypotheses for the mechanism underlying the benefit of cofeeding. To enable parallel, replicated cultures at both metabolic and isotopic steady state, we adapted AneVO, our anaerobic eVOLVER-based mini-reactor platform (Hoyt et al., 2026) into a benchtop anaerobic chemostat capable of maintaining anaerobicity for over 500 hours. The resulting flux maps distinguish between the energetic and flux-redistribution hypotheses, reveal an unexpected oxidative directionality of PFOR, and identify cysteine as a significant and underappreciated carbon source in acetogenic fermentation.

## 2. Materials and Methods

### 2.1 Chemicals and reagents

Media components were obtained in the highest grade available from Fisher Scientific or Sigma Aldrich. ^13^C stable isotopic tracers, including [3-^13^C]-L-cysteine, ^13^C-methanol, and [U-^13^C]-D-glucose, were obtained from Cambridge Isotope Labs.

### 2.2 Microbial strains, routine culturing and media conditions

Unless otherwise stated, cells were cultivated anaerobically in fully defined basal medium with 3 mM cysteine-HCl as a reducing agent (Kountz et al., 2020), as described previously (Hoyt et al., 2026). Wild-type *Eubacterium limosum* (ATCC 8486) was used for batch experiments in Hungate tubes comparing potential sulfur sources and for tracer experiments with labeled cysteine. Chemostat experiments for ^13^C-MFA used a thiamphenicol-resistant, EPS-deficient strain Δ(*epsABCptkA*)::*catP* generated previously (Sanford et al., 2023) to reduce biofilm formation and minimize contamination in long-term cultures. Batch cultures were inoculated from pre-cultures grown with ^13^C tracers matching the experimental conditions, with a 10-fold dilution.

### 2.3 Benchtop anaerobic chemostat growth in AneVO

All ^13^C-MFA experiments were conducted with cells grown in continuous culture, in our previously described AneVO benchtop anaerobic parallel mini bioreactor platform (Hoyt et al., 2026), modified for chemostat mode. Culture vials contained 25 mL anaerobic media with an initial headspace of 10% CO_2_, 5% H_2_ and 85% N_2_, and were sealed with butyl rubber stoppers and aluminum crimp tops. Fresh anaerobic media was supplied from an inverted 500 mL GL45 media bottle sealed with a DURAN^TM^ bromobutyl septum (DWK Life Sciences #292062803), connected to the culture vials via oxygen-impermeable PharMed tubing, needles and Luer-lock fittings. A high-precision multichannel peristaltic pump (Ismatec IPC High Precision Multichannel Pump) was used to deliver fresh media and remove culture at 8.33 µL minute^-1^. The length of the needle for outflow from each vial was selected to set the working volume to 25 mL, resulting in a dilution rate of 0.02 h^-1^. To avoid problems with pressure drop during liquid feeding, the feed bottle was kept under positive pressure with N_2_, and effluent culture was collected in partially evacuated serum bottles. The feed contained basal media supplemented with 50 mM sodium bicarbonate, and either 2.3 mM glucose, 50 mM methanol, or both methanol and glucose, as required. A single feed bottle was used for each substrate condition to ensure identical media composition across replicates. The tubing and needles were assembled, autoclaved, and then purged for 1.5 tubing volumes using sterile anaerobic media. Liquid level was verified each day, and waste bottles were changed each day.

### 2.4 Extracellular metabolite quantification

Substrates and extracellular products were quantified in supernatant samples, prepared by centrifugation of culture samples at 20,000g for 20 minutes. Methanol concentration was determined spectrophotometrically by Purpald assay after enzymatic conversion to formaldehyde, as described previously (Hoyt et al., 2026). Glucose, acetate and butyrate were determined by HPLC-RID (Agilent 1260) using an ion exchange column (BioRad Aminex HPX-87H, 7.8 x 300mm) operated at 60°C, with isocratic elution with 14 mM sulfuric acid at 0.6 mL min^-1^ (Hoyt et al., 2026).

Cysteine concentration was measured after monobromobimane (mBBr) derivatization using an HPLC-FLD (Agilent 1260) equipped with a C18 column (Agilent InfinityLab Poroshell, 120 EC-C18, 2.7 μm 3×100 mm), as described previously (Hayes et al., 2023). Briefly, 10 µL sample was reduced by mixing with 2 µL 1-octanol and 3 µL reducing solution (9M NaBH_4_ in 0.1N NaOH and DMSO at a 2:1 volumetric ratio) for 15 minutes, then quenched with 0.6 µL 3N HCl. 90 µL of mBBr solution (3 mM mBBr in 0.2 mM EDTA, 200 mM Tris, pH 9.5) was added, and the derivatization allowed to proceed at room temperature for 2 hours in the dark, followed by neutralization with 30 µL 1 N HCl. Derivatized samples were centrifuged at 20,000 g for 5 minutes, then transferred into HPLC vials. Mobile phase A was 89.75% Water, 10% Methanol, and 0.25% glacial acetic acid (pH 3.9). Mobile phase B was 89.75% methanol, 10% water, and 0.25% glacial acetic acid (pH 3.9). The flow rate was 0.5 mL/min with gradient held at 0% B from 0 to 2 minutes; ramped to 46% B from 2 to 8 minutes; ramped to 64% B from 8 to 9 minutes; ramped to 100% B from 9 to 13 minutes; ramped to 0% B from 13 to 14 minutes, held at 0% B from 14-15 minutes. The injection volume was 10 µL, column temperature was 40°C, and the excitation/emission wavelengths was 384/478 nm. Quantification was achieved by comparison of peak areas to authentic standards derivatized by the same procedure.

Specific substrate uptake rates and product secretion rates were calculated using the measured inlet substrate concentration and the concentration in the culture as determined above, along with the steady state biomass concentration determined by spectrophotometer OD_600_ measurements and an OD:DCW ratio of 0.29 gDCW/L per OD-L (Loubiere et al., 1992).

### 2.5 Isotope analysis in proteinogenic amino acids, acetate and butyrate

Isotopologue distribution was determined by LC-HRMS, using hydrolyzed biomass samples for proteinogenic amino acids, and derivatized supernatants for acetate and butyrate, as described previously (Hoyt et al., 2026). Raw isotopic abundance was determined using Agilent’s Mass Hunter Profinder software. For derivatized products, the raw isotopic abundances were corrected for the natural ^13^C abundance of the 2-NPH derivatizing agent using Isocor (Millard et al., 2019).

### 2.6 Core metabolic model for *Eubacterium limosum*

The goal of this work was to determine net and exchange fluxes through central metabolic pathways, to understand metabolic differences between single-substrate and cofeeding. For this, we generated a core metabolic model for *E. limosum*, using published biochemistry and genomic expression data for *E. limosum* (Dietrich et al., 2021; Shin et al., 2023; Song et al., 2020, 2018, 2017; Wood et al., 2023). As *E. limosum* is closely related to another methylotrophic acetogen, *Eubacterium callanderi*, we also drew on published biochemical information for this organism (Feng et al., 2022; Flaiz et al., 2024; Poehlein et al., 2024). Finally, ^13^C-MFA has previously been conducted with the acetogen *Clostridium ljungdahlii* (Dahle et al., 2022) and the solventogen *Clostridium acetobutylicum* (Amador-Noguez et al., 2010; Au et al., 2014), which provided helpful starting points.

The full model is shown in the Supplementary Material. Key organism-specific details and assumptions are as follows: 1) Acetogenic methanol metabolism requires a reversal of flux through the methyl branch of the Wood-Ljungdahl pathway to generate reducing equivalents, in addition to a corrinoid methyltransferase to transfer the methyl group from methanol to tetrahydrofolate (Dietrich et al., 2021). 2) Butyrate production is coupled to energy conservation through the bifurcating butyryl-CoA dehydrogenase (encoded by *bcd-etfAB (B2M23_RS01165-1175)*). 3) *E. limosum* does not encode the glycine synthase reductase pathway (Song et al., 2020), the oxidative pentose phosphate pathway or the Entner-Doudoroff pathway (Song et al., 2018). 4) Initial amino acid biosynthesis pathways were taken from previous *C. acetobutylicum* (Au et al., 2014) and *C. ljungdahlii* (Dahle et al., 2022) models, as amino acid biosynthesis is well conserved across acetogens (Shin et al., 2016). 5) PFOR was modelled as two reversible steps: one for decarboxylation of pyruvate, and one for reduction of ferredoxin. This approach was first established for *C. acetobutylicum* to account for differential reversibility between the decarboxylation and reduction reactions (Amador-Noguez et al., 2010). 6) Like other Clostridia, *E. limosum* encodes a bifurcated TCA cycle (Amador-Noguez et al., 2010; Au et al., 2014), and lacks succinate dehydrogenase (Song et al., 2018). 7) Cysteine uptake and metabolism through cysteine desulfurase were included (see discussion in Section 3.1). 8) The Rnf complex was included in the model as a reversible reaction with the assumption that the reduction of NAD^+^ by Fd_red_ is coupled to the export of 2 Na^+^, which are subsequently imported by the ATP synthase to generate 0.5 ATP (Gasparrini et al., 2025). 9) The physiological electron donor for methylene-THF reductase (MTHFR) in *E. limosum* is not known. Here we assume an electron-bifurcating MTHFR (Kremp and Müller, 2021), in which electrons from NADH are bifurcated to methylene-THF and Fd_ox_. 10) Finally, as for other Clostridia, we include a CO_2_ exchange reaction to account for dilution of intracellular ^13^CO_2_ by uncharacterized unlabeled CO_2_ sources (Au et al., 2014).

### 2.7 ^13^C-metabolic flux analysis and ΔG calculation

Steady-state fluxes were calculated using ^13^C mass isotopologue distributions (MIDs) obtained from proteinogenic amino acids and extracellular products, as well as the measured specific uptake and production rates, and the growth rate (0.02 h^-1^). Net CO_2_ production or consumption flux was not measured directly, and calculated during MFA. The flux estimations and confidence intervals were generated using Openflux2 (Quek et al., 2009; Shupletsov et al., 2014). The MID inputs, as well as the uptake and productivity basis measurements, were averages of triplicate biological replicates, except for the glucose/CO_2_ model which was in quadruplicate, and the standard deviation was used as the error for flux fitting. Flux estimation was conducted by minimizing the sum-squared residuals between predicted and measured MIDs, starting from 500 randomized initial flux estimates to ensure identification of the best global solution. Goodness-of-fit was determined by chi-square test, and 95% confidence intervals were obtained using nonlinear sensitivity analysis for individual net fluxes and reversibility ratios (Quek et al., 2009; Shupletsov et al., 2014). The Gibbs free energy for PFOR was calculated directly from the flux reversibility ratio, according to (Park et al., 2016):

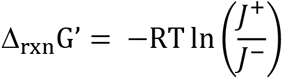

where R is the universal gas constant, T is reaction temperature, and 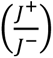 is the reversibility ratio. All fluxes, including upper and lower bounds and statistical analysis, are reported in the Supplementary Material. The model input file, including the stoichiometry and carbon transitions is shown in Table S7, MID values are in Table S8, and the flux distributions are in Table S9. All presented flux units are scaled by a factor of 100 to reduce decimal truncation during fitting (e.g. 200 represents 2 mmol/gDCW-h).

### 2.8 Cofactor balances

Energy cofactor generation and consumption (NADH, NADPH, Fd_red_ and ATP) were balanced for the base model during MFA, and the net fluxes determined were used to compare the energy cofactor contributions from each pathway (Dietrich et al., 2021). To evaluate an alternative scenario where MTHFR accepts electrons from reduced ferredoxin through the RNF complex in an energy-conserving manner (Öppinger et al., 2022), carbon fluxes from the base model were used, with the revised stoichiometry for MTHFR/RNF complex of methylene-THF + Fd_red_ + 0.5 ADP ↔ methyl-THF + Fd_ox_ + 0.5 ATP. The NADH and reduced ferredoxin were balanced post-hoc by manually setting the standalone RNF flux. In all cases, excess ATP was ascribed to maintenance energy (Dahle et al., 2022). Cofactors ascribed to ‘Biomass + AA’ included both the direct contribution from the biomass equation, and amino acid biosynthetic reactions including the bifurcated TCA.

### 2.9 Differential gene expression analysis

RNA was isolated from chemostat cultures by commercial kit (NEB, Monarch Total RNA Miniprep kit), and submitted for library generation, sequencing and analysis by commercial vendor (Azenta). Culture samples were collected anaerobically from the chemostat during steady state growth, preserved in equal volume of RNA Protect (New England Biolabs, NEB), centrifuged, and the cell pellets were stored at -80°C until use. On the day of RNA extraction, cells were resuspended in nuclease free TE buffer (IDT) supplemented with 1 mg/mL lysozyme for lysis via bead beating. 250 µL of resuspended cells were added to commercially prepared 2-mL tubes containing acid washed glass beads (Thomas Scientific) and then lysed by horizontal vortex (Scientific Industries Vortex Genie 2) for 15 minutes (Sanford and Woolston, 2022b). 500 µL of RNA lysis buffer (NEB) was added to the tube, and vortexed for 10 seconds. Lysate was centrifuged at 16,000g for 10 minutes. RNA extraction was conducted using the Monarch Total RNA Miniprep kit (NEB), as prescribed by the kit protocol, including DNAse cleanup treatment. The approximate concentration of extracted RNA was confirmed using a NanoDrop One UV– Vis Spectrophotometer (ThermoFisher Scientific). The RNA library preparation and sequencing was performed by Azenta Life Sciences using Illumina 2×150 bp with a target of 10 million reads per sample. Differential gene expression was determined by averaging the gene transcript counts for each growth condition replicate then calculating the log_2_-fold change.

## 3. Results and Discussion

### 3.1 *E. limosum* Incorporates Cysteine into Central Metabolism

^13^C-MFA requires a defined medium to accurately account for the uptake of all carbon sources (Krömer et al., 2014; Long and Antoniewicz, 2019). A fully defined medium has been established for *E. limosum* (Hoyt et al., 2026), but includes cysteine as a reducing agent to ensure a sufficiently low redox potential for anaerobic growth (Mariotto et al., 1989). Cysteine can also serve as a nutrient, as recently observed in *Moorella thermoacetica*, supplying organic sulfur, electrons, carbon, and nitrogen (Göbbels et al., 2021). To test whether cysteine uptake could confound flux estimations, we grew *E. limosum* on glucose and measured the cysteine concentration over time (**Figure 1**). Cysteine concentration decreased over time at approximately 0.1 mmol/gDCW-h (**Figure 1a**). We therefore initially attempted to eliminate cysteine from the media. Other reducing agents and sulfur sources have been reported for *E. limosum,* including sodium sulfide (Mariotto et al., 1989), as well as thiosulfate after adaptive laboratory evolution (Pregnon et al., 2022). However, in our hands, growth with either of these compounds was poor, exhibiting long lag times and low cell densities compared to growth with cysteine (**Supplementary Table S1**).

**Figure 1.**
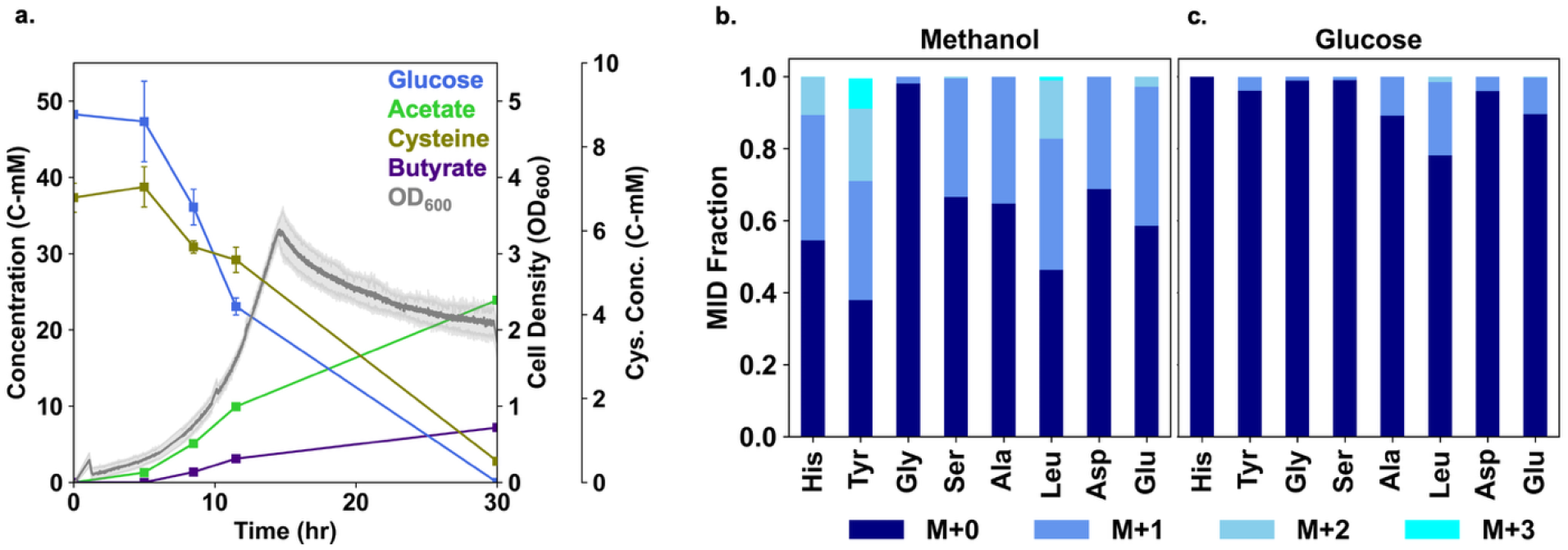
Cysteine is consumed during growth on either methanol or glucose. a) Growth plot for *E. limosum* grown on glucose in defined medium supplemented with cysteine. Mass Isotopologue Distributions (MID) for proteinogenic amino acids obtained from *E. limosum* grown with [3-_13_C] cysteine, with methanol (b) or glucose (c). Standard deviation for MID measurements were less than 1% for glucose-grown cells, and less than 6% for methanol-grown cells.

Instead, since the benefit of methanol-glucose cofeeding is observed in the presence of cysteine (Hoyt et al., 2026; Loubiere et al., 1992), and its concentration in culture can be measured, we elected to leave the media unchanged and account for the uptake of cysteine in the MFA. This required determining by what pathway cysteine is metabolized, the impacts to carbon backbone transitions, and updating the metabolic model accordingly. Both cysteine desulfurase and cysteine desulfidase incorporate cysteine into pyruvate with no changes to the carbon backbone. *E. limosum* has several annotated cysteine desulfurases [EC 2.8.1.7], with one of them – encoded by B2M23_RS07680 – expressed highly under both autotrophic and heterotrophic conditions, but no annotated desulfidases (Song et al., 2017). Cysteine desulfurase catalyzes the conversion of cysteine into alanine, releasing a sulfur atom that is either transferred to an iron-sulfur cluster or released as H_2_S. Alanine then enters central metabolism through conversion to pyruvate by a transaminase. To confirm this route, we supplemented *E. limosum* with [3-^13^C] cysteine and analyzed the labeling of proteinogenic amino acids during growth on either methanol (**Fig. 1b**) or glucose (**Fig. 1c, Figure S2)**. Consistent with the activity of cysteine desulfurase, a substantial fraction of the alanine pool was M+1 labeled. Although labeling was observed under both growth conditions, the fraction of labeling was significantly higher during growth on methanol. Indeed, the labelled fractions total ∼50% for tyrosine, histidine and leucine, with substantial M+2 labeling. Tyrosine and histidine are derived from intermediates of the pentose phosphate pathway, where carbon rearrangements involving G3P and F6P generate doubly labeled compounds. The presence of doubly labeled leucine is consistent with its biosynthesis from two molecules of pyruvate. Collectively, these data indicate a significant flux of carbon from cysteine to pyruvate, most likely via cysteine desulfurase, which cannot be ignored in ^13^C-MFA.

### 3.2 AneVO chemostat culture enables isotopic and metabolic stationarity and exhibits the benefit of cofeeding

Our prior work on glucose/methanol cofeeding in *E. limosum* was conducted in fed-batch culture, where methanol was present in the medium, and glucose was dosed slowly over time. ^13^C tracer experiments in this configuration showed significant variability in isotopic labeling at different time points during the growth phase (Hoyt et al., 2026). Since metabolic and isotopic stationarity are a prerequisite for steady-state ^13^C-MFA, this made determining quantitative fluxes impossible. To solve this problem, here we reconfigured AneVO – the parallel benchtop anaerobic mini bioreactor system we previously developed – for chemostat growth. Briefly, this involved using a high-precision multichannel peristaltic pump with narrow-bore oxygen-impermeable tubing to enable sufficiently low flow in and out of each culture vial for the desired dilution rate, and using N_2_-pressurized feed bottles and partially evacuated serum bottles for waste collection to ensure flow consistency. The full setup is shown in **Supplementary Figure S1**.

Pilot tests in the AneVO chemostat with unlabeled substrates identified that a dilution rate of 0.02 h^-1^ and methanol-to-glucose feed ratio of 25 to 1 led to simultaneous consumption of both methanol and glucose (**Figure S3**). Importantly, these conditions still resulted in a growth benefit from glucose cofeeding compared to methanol-only controls, with a steady state OD_600_ approximately 3-fold higher. With these conditions identified, we repeated the chemostat experiments with ^13^C tracers for flux analysis (Methanol: ^13^C-methanol; Glucose: U-^13^C-glucose; Cofeeding: ^13^C-methanol + unlabeled glucose). All conditions contained unlabeled CO_2_ in the form of bicarbonate in the medium.

To assess metabolic and isotopic stationarity, samples were collected daily and analyzed for substrate and product concentrations, cell density, and ^13^C incorporation (**Figure 2, Tables S2, S3, S4**). All growth conditions resulted in a steady isotopic state (assessed by MIE data) that lasted at least 100 hours (**Fig. 2a-c**). Notably, the standard deviations of all MIEs across the timepoints during steady state were less than 1%. By comparison, in our previous fed-batch work, amino acid labeling changed by an average of 16% between different time points during the growth phase. This highlights the benefit of continuous culture for rigorous ^13^C-MFA. Within the steady state time window, cell density and product concentrations were remarkably stable (**Fig. 2d-f**), particularly for co-fed and glucose-grown cells. Methanol-grown remained stable for ∼70 hours, after which the three replicate vials began to diverge (**Fig. S4a-c**), resulting in varied cell density and butyrate concentration. The measurements collected when the three replicates were closest in growth (426 hours, red arrow in **Fig. 2e**) were used for flux analysis. Compared to the other substrates and products, methanol measurements were more variable. We suspect this reflects measurement error due to challenges associated with the volatility of methanol or imprecision in the enzymatic assay, rather than actual variation in methanol uptake rate, because the MIE and product concentrations show very little change during the steady state period. To account for this variability, the methanol concentration was averaged across the five time points surrounding the one used for labeling data for MFA.

**Figure 2.**
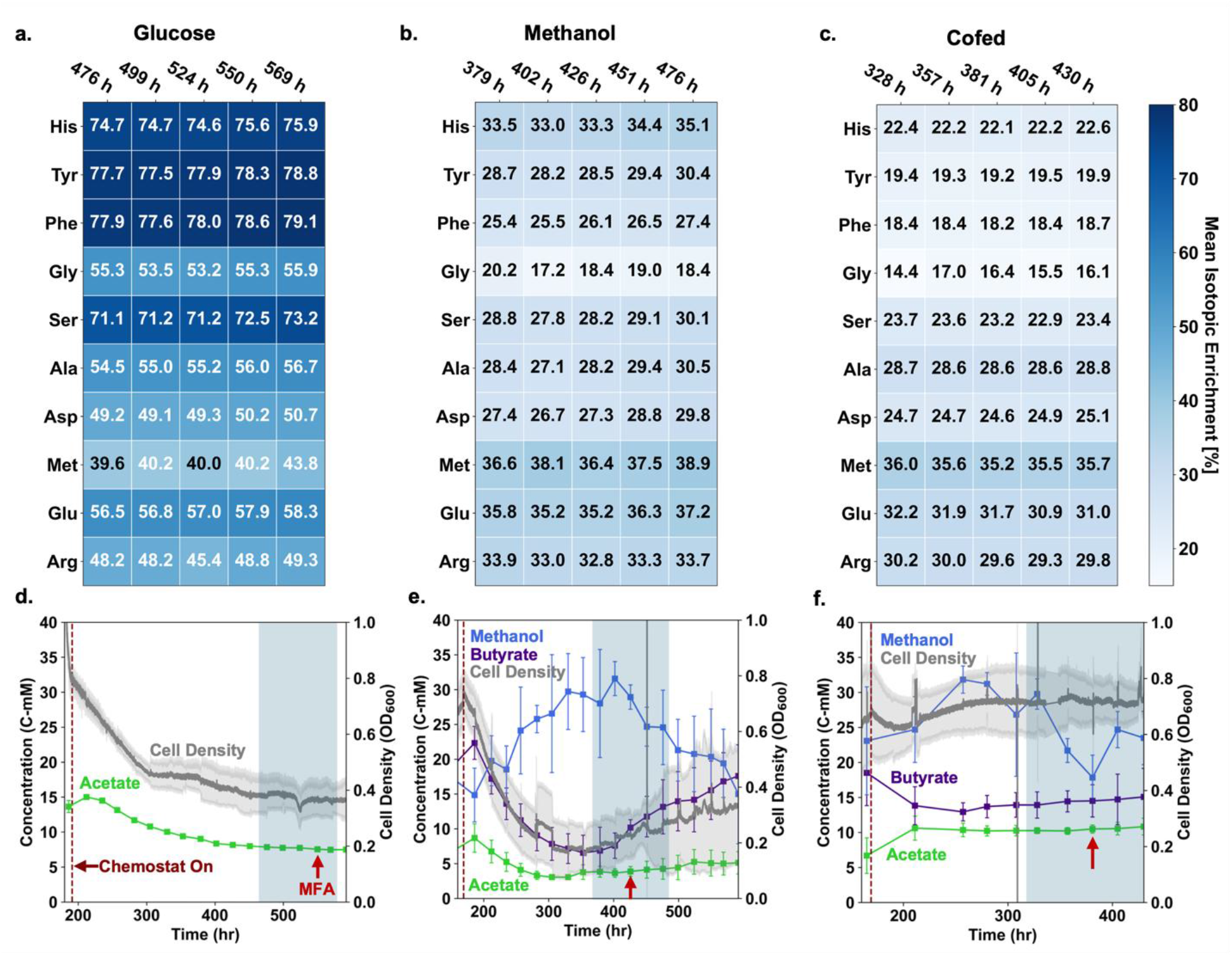
Chemostat experiments reached stationarity in the AneVO chemostat. Heat maps show Mass Isotopic Enrichment (MIE) values for each substrate condition: a) glucose/CO_2_ grown, b) methanol/CO_2_, c) cofed. The MIE standard deviations were 5% or less across growth replicates for the data shown, and were 2.5% or less for timepoints used for MFA. All times are provided as cumulative values, with methanol inoculations occurring at t=0 hours, and glucose inoculations occurring at t=168h. Growth data are shown for each substrate condition: d) glucose/CO_2_ grown, e) methanol/CO_2_, f) cofed. The shading identifies the time period shown above in the heat map. The beginning of chemostat operation is indicated with the vertical dashed line. Red arrows show time point used for MFA. For the glucose/CO_2_ substrate condition, chemostat operation began at t=191 hours and measurements at t=550 hours were used for MFA. For the methanol/CO_2_ substrate condition, chemostat operation began at t=168 hours and measurements for time point t=426 hours were used right before cultures diverged. For the cofed substrate condition, chemostat operation began at t=169 hours and measurements from t=381 hours were used for MFA. Glucose was fully consumed and did not accumulate in the media, per HPLC measurements.

The substrate, product and biomass concentrations obtained from continuous culture were next used to calculate steady state specific uptake and production rates, and biomass yields (**Table 1**). Methanol was consumed during both methylotrophic growth and cofeeding. Glucose was undetectable in cultures grown either heterotrophically or under cofeeding conditions. These results confirm simultaneous substrate utilization in the cofeeding experiments, suggesting that glucose limitation in continuous culture avoided problems with catabolite repression previously reported in batch cultures, where methanol uptake was inhibited by the presence of glucose (Bae et al., 2026; Hoyt et al., 2026; Loubiere et al., 1992).

**Table 1.**
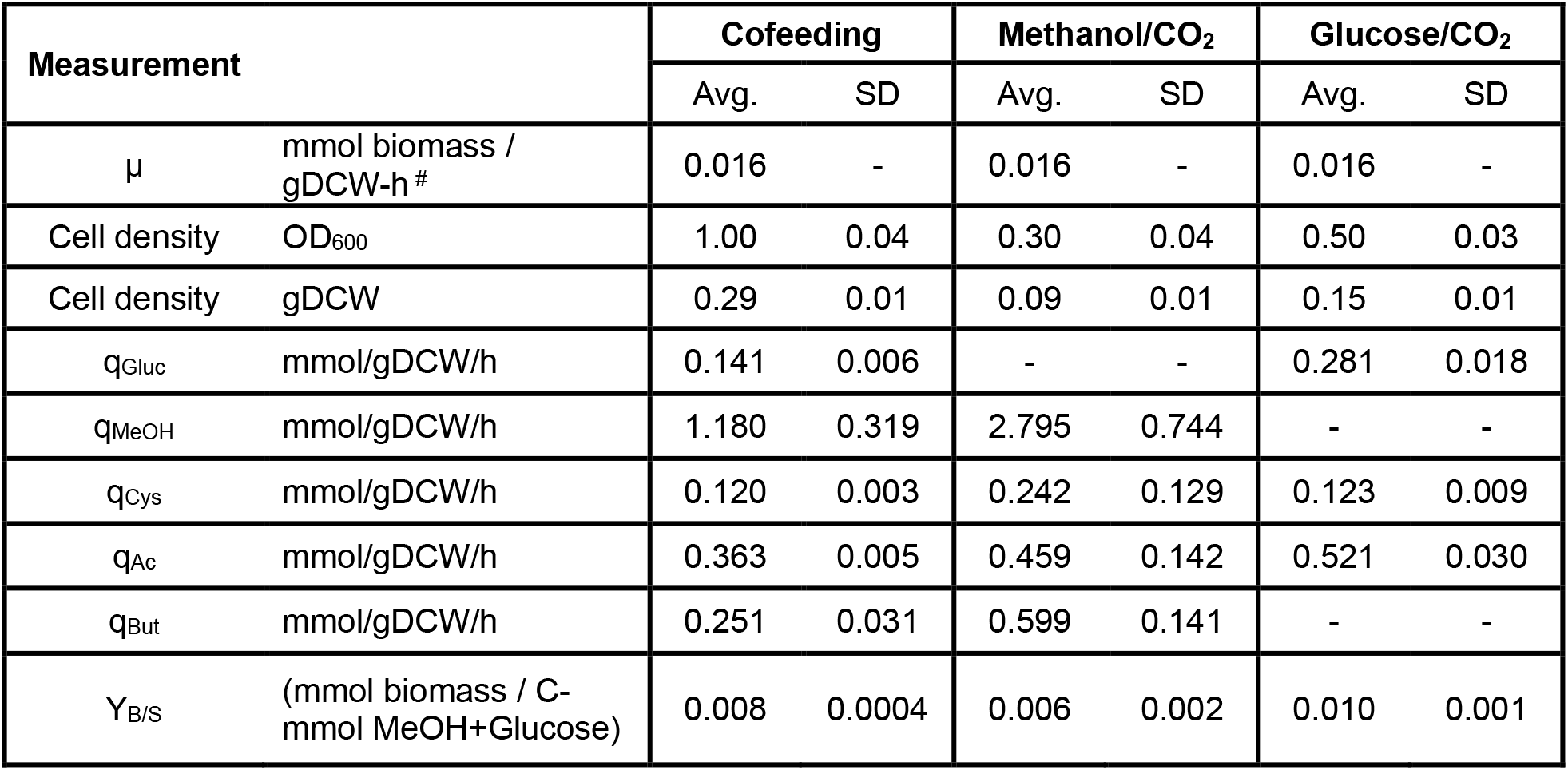
MFA basis calculations from chemostat growth experiments. ^#^Molar specific growth rate calculated by dividing the growth rate of 0.02 h^-1^ by the molecular weight of biomass, then multiplying by 1,000 mmol/mol. The molecular weight of biomass used was 1244.15 g/mol (Au et al., 2014), based on the stoichiometric coefficients used in the biomass equation, used in the model.

| Measurement | | Cofeeding | | Methanol/$\text{CO}_2$ | | Glucose/$\text{CO}_2$ | |
| --- | --- | --- | --- | --- | --- | --- | --- |
|  |  | Avg. | SD | Avg. | SD | Avg. | SD |
| $\mu$ | mmol biomass / gDCW-h<sup>#</sup> | 0.016 | - | 0.016 | - | 0.016 | - |
| Cell density | OD<sub>600</sub> | 1.00 | 0.04 | 0.30 | 0.04 | 0.50 | 0.03 |
| Cell density | gDCW | 0.29 | 0.01 | 0.09 | 0.01 | 0.15 | 0.01 |
| $q_{\text{Gluc}}$ | mmol/gDCW/h | 0.141 | 0.006 | - | - | 0.281 | 0.018 |
| $q_{\text{MeOH}}$ | mmol/gDCW/h | 1.180 | 0.319 | 2.795 | 0.744 | - | - |
| $q_{\text{Cys}}$ | mmol/gDCW/h | 0.120 | 0.003 | 0.242 | 0.129 | 0.123 | 0.009 |
| $q_{\text{Ac}}$ | mmol/gDCW/h | 0.363 | 0.005 | 0.459 | 0.142 | 0.521 | 0.030 |
| $q_{\text{But}}$ | mmol/gDCW/h | 0.251 | 0.031 | 0.599 | 0.141 | - | - |
| $Y_{\text{B/S}}$ | (mmol biomass / C-mmol MeOH+Glucose) | 0.008 | 0.0004 | 0.006 | 0.002 | 0.010 | 0.001 |

Comparing between culture conditions, cofeeding resulted in lower specific uptake rates of each substrate (1.180 mmol/gDCW-h vs 2.795 mmol/gDCW-h for methanol, and 0.141 mmol/gDCW-h vs. 0.281 mmol/gDCW-h for glucose), yet the highest cell densities (OD 1.00 ± 0.04, vs 0.30 ± 0.04 for methanol and 0.5 ± 0.03 for glucose, measured by spectrophotometer). Specific uptake rates of cysteine were statistically indistinguishable across culture conditions. The biomass yield (Y_X/S_) – calculated on the basis of total C-mol in glucose and methanol – was ∼33% higher under cofeeding conditions than on methanol alone. This indicates that the benefit of cofeeding leads to increased metabolic efficiency over methanol alone, rather than simply an increase in total carbon availability.

Butyrate production was detected only in cultures supplemented with methanol, which was expected (Litty and Müller, 2021). Both acetate and butyrate titers were higher during cofeeding than under other conditions, although the specific productivity for both products was highest during growth on methanol, and the butyrate:acetate production ratio was also lower during cofeeding. Overall, these experiments validate AneVO for continuous anaerobic growth of *E. limosum,* confirm a metabolic benefit of cofeeding, and provide a temporally consistent isotopic dataset of much higher quality than fed-batch cultivation, suitable for mechanistic comparison of cofeeding and single-substrate conditions by ^13^C-MFA.

### 3.3 The Wood-Ljungdahl pathway is active during heterotrophic growth

The specific substrate uptake and production rates, along with the isotopic data from proteinogenic amino acids, acetate and butyrate, were used to fit fluxes under each condition. All three fits were statistically acceptable, with the sum of the residuals well below the 95% chi-squared threshold (**Table S5**), for the full model shown in **Table S6** and the MID fits shown in **Table S7**. The carbon and electron balances derived from simulated fluxes closed to within 92% (**Table S8).** Additionally, confidence intervals around all net fluxes were fully resolved (**Table S9**), as was the flux ratio (J^+^/J^-^) for the PFOR-catalyzed reaction (**Table S10, Table S11**). Collectively, these results validate the core metabolic model for *Eubacterium limosum*.

We first consider growth on glucose, for which the flux distribution is shown in **Figure 3**. Acetate is the primary fermentation product under these conditions, with a yield of 2.4 mol acetate per mol glucose. The maximum theoretical yield for homoacetogenesis, where reducing equivalents derived from the conversion of glucose to two mol of acetyl-CoA through glycolysis and PFOR are used to fix an additional mol of acetyl-CoA through the WLP, is 3. The slightly lower value here reflects carbon diverted to biomass, and a small amount of net CO_2_ production (0.06 mmol/gDCW-h), as in previous work with *Moorella thermoacetica* (Park et al., 2019). Consistent with this yield and the WLP functioning as an electron sink under heterotrophic conditions, ^13^C-MFA revealed that 30% of the acetyl-CoA is derived from CO_2_ fixation. These results reinforce the essentiality of the WLP during growth on sugars. In a recent genome-wide CRISPRi study, downregulation of WLP genes resulted in a significant growth defect (Shin et al., 2023).

**Figure 3.**
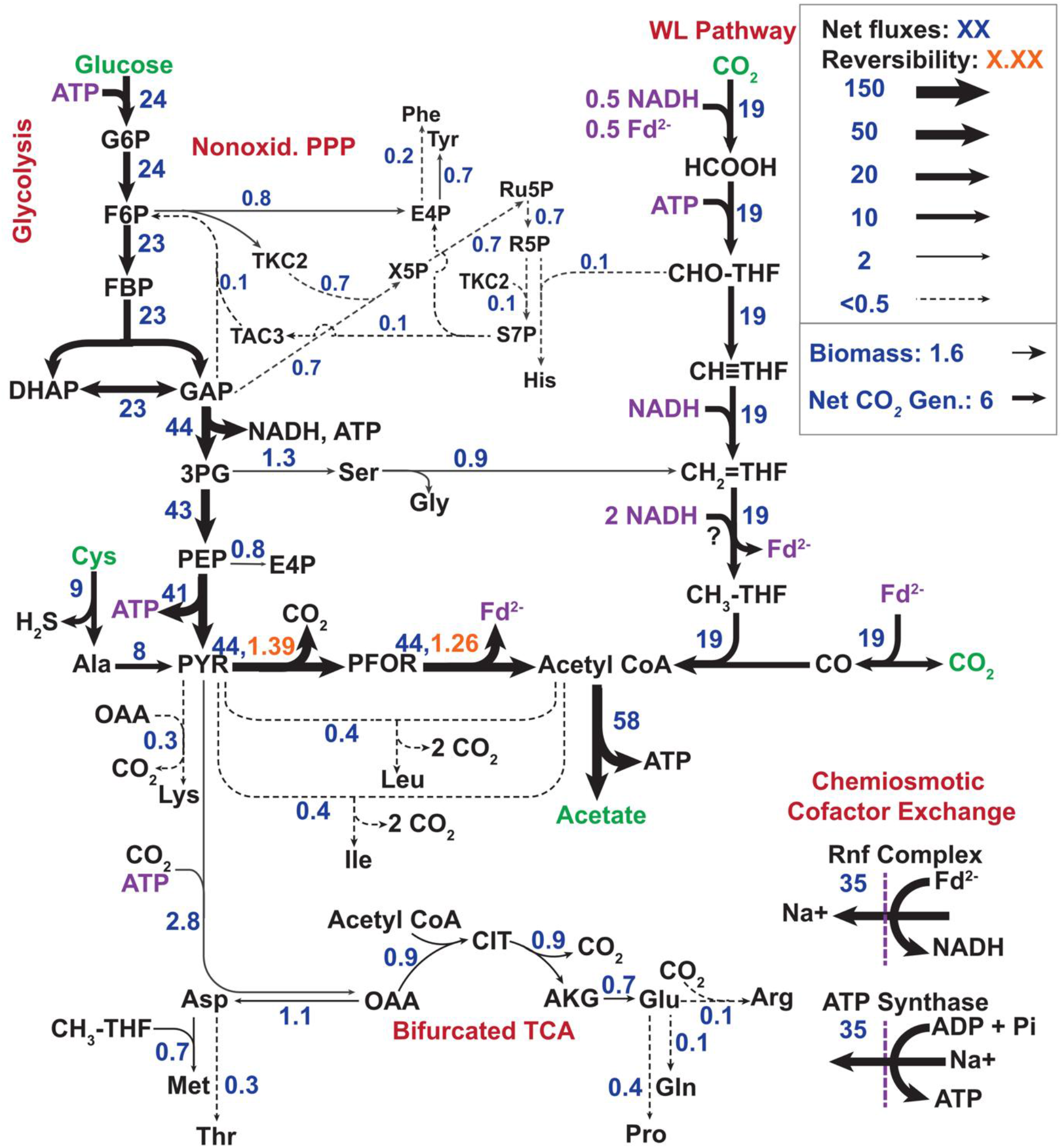
^13^C MFA Carbon Flux Distribution for *E. limosum* grown on Glucose/CO_2_. For readability and to reduce decimal truncation, fluxes are scaled by a factor of 100; a flux of 100 = 1 mmol/gDCW-h. Line thicknesses correspond to natural log of approximate fluxes including the scaling factor. Net fluxes are shown with 95% confidence intervals presented in **Table S9**. Key pathway names used in the text are shown in red. Substrates and products are shown in green. Flux values are based on reaction substrate, and shown in blue.

There is also very little cross-talk between glycolysis and the WLP, with <5% of the flux to methylene-THF coming from serine hydroxymethyltransferase. This is consistent with the low ^13^C enrichment of the methyl group in methyl-THF (8%) derived from isotope balance analysis of the relative labeling of methionine and aspartate (**Table S12).** Finally, as expected during heterotrophic growth, flux through PFOR was in the oxidative direction. Interestingly, approximately 16% of the flux to pyruvate was from cysteine, resulting in substantial fractions of completely unlabeled (M+0) amino acids derived from glycolytic intermediates (Alanine, 26.2%; Serine, 16%; Glycine, 27.8%; **Table S8**) and highlighting the importance of accounting for cysteine uptake.

### 3.4 ^13^C-MFA reveals oxidative flux through PFOR and reverse flux through Rnf during methylotrophic growth

The fluxes for *E. limosum* grown on ^13^C methanol and unlabeled CO_2_ are shown in **Figure 4**. Under these conditions, butyrate and acetate are produced at similar rates (0.60 and 0.46 mmol/gDCW-h, respectively), due to the extra reducing equivalents in methanol compared to glucose. In general, the measured fluxes agree with previous observations and proposals for acetogenic methanol conversion (Kremp and Müller, 2021): For every mol of methanol that enters the methyl branch as methyl-THF, 0.4 are oxidized to produce reducing equivalents, while 0.6 are directly incorporated through ACS/CODH to produce acetyl-CoA. The molar methanol:CO_2_ uptake ratio is 4.7:1, highlighting the unique ability of acetogens for net fixation of CO_2_ during methylotrophic growth (Gasparrini et al., 2025; Sanford and Woolston, 2022a). Oxidation of methanol through the methyl branch of the WLP provides primarily NADH. Reduced ferredoxin needed for conversion of CO_2_ to CO in the carbonyl branch is predicted to come from reversing electron flow through the Rnf complex, using the sodium motive force to couple NADH oxidation to Fd reduction at the expense of ATP (Dietrich et al., 2021). Indeed, this flux is one of the largest in the network (1.4 mmol/gDCW-h) during growth on methanol, second only to CODH/ACS.

**Figure 4.**
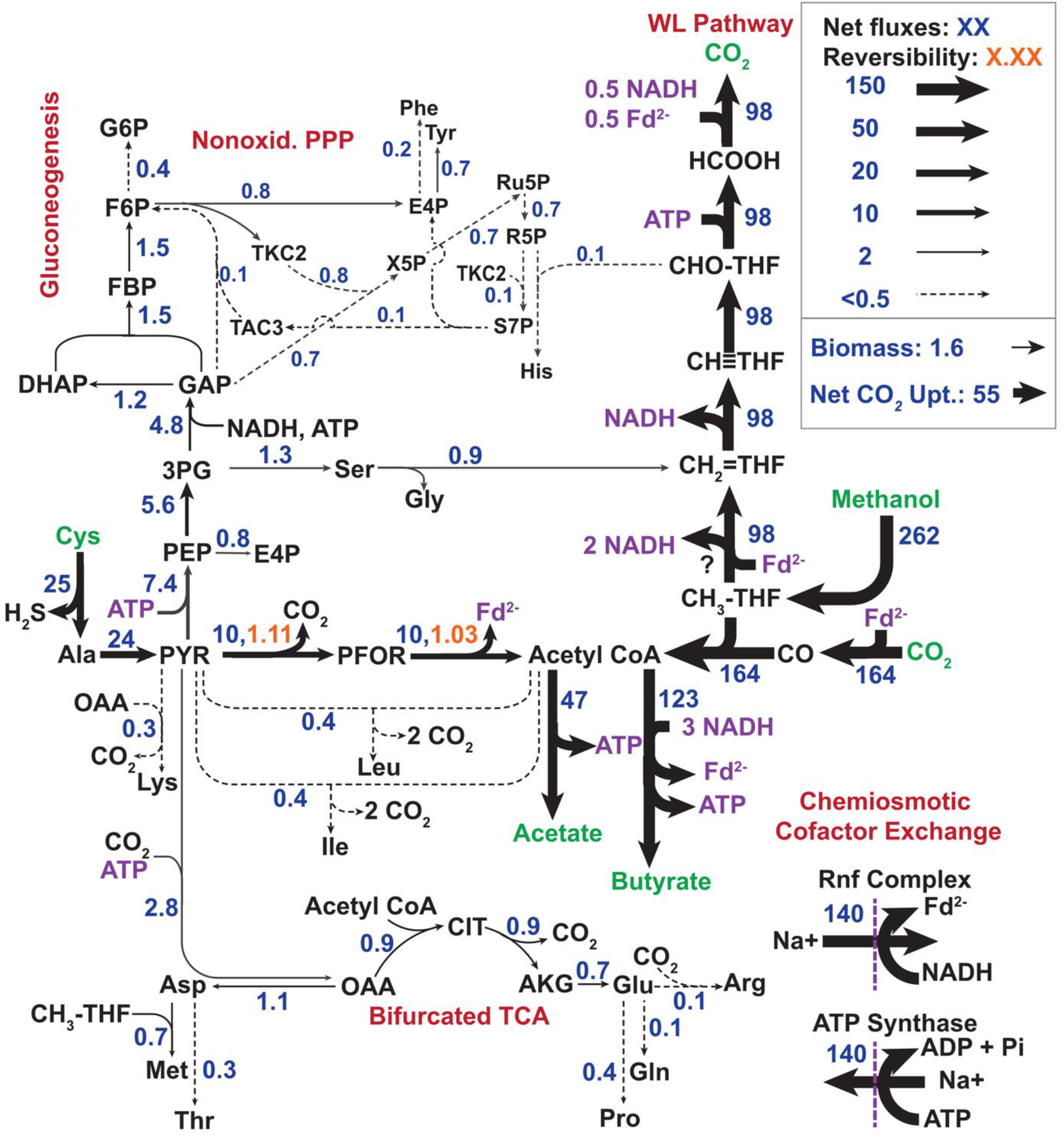
^13^C MFA Carbon Flux Distribution for *E. limosum* grown on methanol/CO_2_. For readability and to reduce decimal truncation, fluxes are scaled by a factor of 100; a flux of 100 = 1 mmol/gDCW-h. Line thicknesses correspond to natural log of approximate fluxes including the scaling factor. Net fluxes are shown with 95% confidence intervals presented in **Table S9**. Key pathway names used in the text are shown in red. Substrates and products are shown in green. Flux values are based on reaction substrate, and shown in blue.

Interestingly, the ^13^C data reveal a degree of reversibility in the methyl branch during growth on methanol. Although ^13^C-MFA could not quantitatively resolve the exchange fluxes in this branch, isotope balancing revealed that methyl-THF was only 77% labeled (**Table S12**), which can only result from reverse flux of unlabeled CO_2_. In *E. limosum*, the net flux direction of the methyl branch of the WLP completely reverses depending on the substrate, using the reductive direction for CO_2_ growth, but the oxidative direction for methanol growth (Dietrich et al., 2021). To achieve this flexibility, the methyl branch of *E. limosum* may operate very near to equilibrium, which requires carrying some flux in both directions, and producing high amounts of enzyme to achieve net flux in the needed direction (Noor et al., 2014).

Surprisingly, flux analysis revealed net oxidative flux (from pyruvate to acetyl-CoA) through PFOR during methylotrophy. Classically, during methylotrophic and autotrophic acetogenesis, flux through PFOR is thought to be in the reductive direction, siphoning acetyl-CoA flux from acetate through gluconeogenesis to produce critical biosynthetic intermediates such as amino acids, nucleotides, and cell wall sugars (Furdui and Ragsdale, 2000). Careful analysis of the pyruvate node reveals that all gluconeogenic flux for these intermediates is actually provided by *cysteine,* rather than acetyl-CoA from the WLP. Approximately half of the cysteine flux to pyruvate (0.24 mmol/gDCW-h) is used for biomass precursors (0.07 to gluconeogenesis, 0.012 to other amino acid biosynthesis), with the remainder converted to acetyl-CoA through PFOR (0.10 mmol/gDCW-h). This finding is important for two reasons: First, one of the prevailing hypotheses explaining the improved growth with glucose cofeeding is that cells avoid the need for reductive flux through PFOR, which is thermodynamically challenging (Δ_r_G^’m^ = 20.9 ± 13.1 kJ.mol (Flamholz et al., 2012; Loubiere et al., 1992). The fact that PFOR flux is actually in the oxidative direction during both methylotrophic and heterotrophic growth overturns this hypothesis. Second, it was recently argued from a thermodynamic perspective that the need for reductive PFOR flux during methylotrophic growth in *E. limosum* imposed a limit on the Fd_red_:Fd_ox_ ratio that would make simultaneous reverse electron flow through the Rnf complex (i.e. the oxidation of NADH with Fd_ox_) infeasible (Wood et al., 2023). As shown by our data, the presence of cysteine in the growth media – which was also used in that study – obviates the need for conversion of acetyl-CoA to pyruvate by PFOR, making the proposed reverse flux through Rnf thermodynamically feasible.

### 3.5 PFOR flux is oxidative during cofeeding, and close to equilibrium under all substrate conditions

The flux distribution for *E. limosum* grown under cofeeding conditions (^13^C-methanol and unlabeled glucose) is shown in **Figure 5**. Consistent with both other growth conditions, net flux through PFOR was in the oxidative direction during cofeeding. Thus, the benefit of cofeeding cannot be explained by a reversal in flux (or elimination of flux) through this enzyme, as had been previously hypothesized (Le Bloas et al., 1993; Loubiere et al., 1992). Indeed, net PFOR flux during cofeeding was higher than during methylotrophic growth (0.23 mmol/gDCW-h vs 0.1).

**Figure 5.**
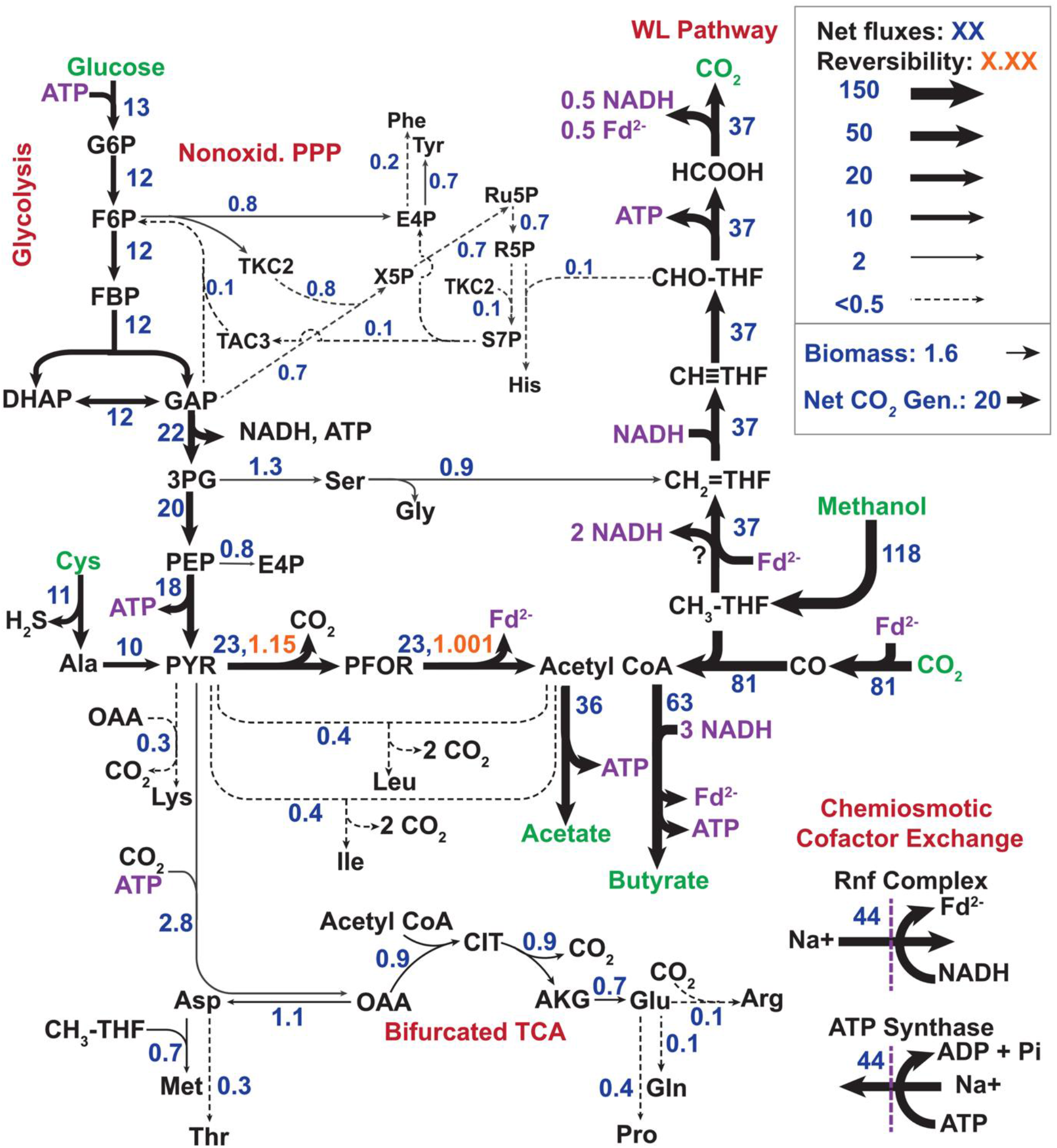
^13^C MFA Carbon Flux Distribution for *E. limosum* cofed with methanol/CO_2_/glucose. For readability and to reduce decimal truncation, fluxes are scaled by a factor of 100; a flux of 100 = 1 mmol/gDCW-h. Line thicknesses correspond to natural log of approximate fluxes including the scaling factor. Net fluxes are shown with 95% confidence intervals presented in **Table S9**. Key pathway names used in the text are shown in red. Substrates and products are shown in green. Flux values are based on reaction substrate, and shown in blue.

To further investigate the flux through PFOR, we used the exchange flux to calculate the ratio of forward:reverse flux (J^+^/J^-^) and *in vivo* ΔG, a powerful feature of ^13^C-MFA (Park et al., 2016). Determining how close a given reaction is to equilibrium allows one to predict whether it is a good target for metabolic engineering to improve overall fluxes. If a reaction is far from equilibrium, flux is kinetically limited, and enzyme overexpression will likely improve pathway flux. If a reaction is close to equilibrium, its rate is primarily constrained by thermodynamics, and changing enzyme concentration will have limited impact on flux (Krömer et al., 2014; Noor et al., 2014; Woolston et al., 2018). Although we were unable to determine confidence intervals for many of the exchange fluxes in *E. limosum* here, we were able to precisely quantify them for PFOR (**Table S10)**, and calculate ΔG (**Table S11)**. Across all substrate conditions, the ratios of forward:reverse flux (J^+^/J^-^) for both the decarboxylation and Fd reduction steps were very close to unity, indicating PFOR is poised very close to equilibrium. Interestingly, both steps are the most forward-driven with glucose (95% CI of J^+^/J^-^ for decarboxylation, 1.389-1.417; for Fd reduction, 1.258-1.313), and closer to equilibrium for growth on methanol (decarboxylation, 1.072-1.142; Fd reduction, 1.001-1.058) and cofeeding (decarboxylation, 1.133-1.155; Fd reduction, 1.001-1.024), with the methanol and cofeeding conditions being nearly indistinguishable. The near-equilibrium operation of PFOR may explain why it is produced so abundantly by acetogens: The closer a reaction to equilibrium, the exponentially more of that enzyme required to sustain a given flux (Noor et al., 2014).

### 3.6 Glucose cofeeding improves growth by avoiding resource-intensive steps in methanol metabolism

Having ruled out changes in PFOR flux as the mechanism underlying enhanced yield during cofeeding, we used the ^13^C-MFA results to probe other possibilities. Chief among these is the hypothesis that pairing a sugar substrate with a C1 substrate increases ATP production: Whereas methylotrophic growth provides ∼0.5 ATP per acetate formed, homoacetogenic acetate production from glucose results in ∼1.5 ATP per acetate, resulting in faster growth and higher biomass yield (Fast et al., 2015; Lee et al., 2022; Liu et al., 2020; Neuendorf et al., 2021; Park et al., 2019). To explore this possibility, the flux distributions were used to calculate the energy and redox cofactor balances for each substrate condition, attributing their production and consumption to specific metabolic pathways as shown in **Figure 6**, with calculations shown in **Supplementary Table S13**. Interestingly, during cofeeding, conversion of glucose to acetate and butyrate only accounts for 20% of the total ATP generation (**Fig. 6a**). Additionally, during methylotrophic growth, reverse flux through the methyl branch of the WLP provides almost as much ATP (through the reversal of formate-tetrahydrofolate ligase) as is generated during cofeeding. This would suggest that glucose cofeeding does not drastically heighten the rate of ATP production.

**Figure 6.**
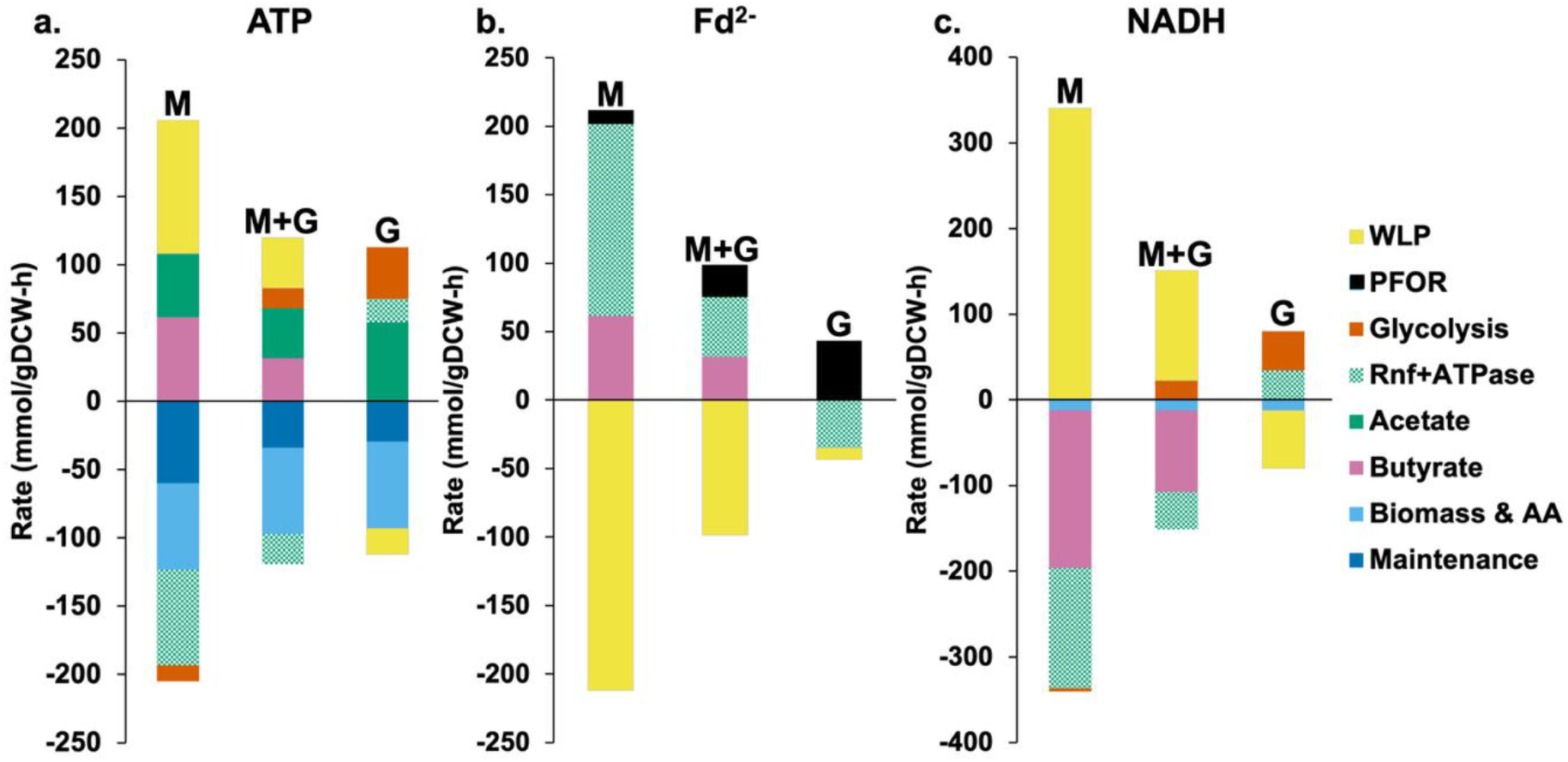
Energy and Redox cofactor balances. Positive values show generation and negative values show consumption. The net generation of each cofactor was calculated from flux distribution and stoichiometric coefficients, and the units shown are consistent with flux maps (100 = 1 mmol/gDCW-h). Flux through the Rnf and ATP synthase chemiosmotic energy balance system was set using a yield 0.5 ATP per 1 NADH per 1 reduced ferredoxin.

However, growth on methanol does indeed require a higher total generation rate of ATP than other conditions (2.05 mmol/gDCW-h vs. ∼1.1 mmol/gDCW-h). The predominant ATP sink is the reverse electron flow through the Rnf complex to produce Fd_red_, which is driven by ATP hydrolysis. This flux is ∼3-fold higher during growth on methanol, due to a significant drop in the specific methanol uptake rate during cofeeding (**Figure 5**, **Table 1**). The other major ATP sink is the maintenance energy, which is >2-fold higher during methylotrophic growth than other conditions. Thus, although glucose contributes little *additional* ATP, the relevant difference between conditions is not how much ATP glucose supplies, but how much the methanol-driven WLP *demands.* Taken together, these data suggest an alternative hypothesis: that methylotrophic WLP is highly resource-intensive, requiring a significant investment of ATP to produce Fd_red_ and for maintenance; and by reducing methanol flux, cofeeding alleviates this heightened demand. Despite the reduced methanol uptake rate, the improved metabolic efficiency of cofeeding nevertheless results in a superior growth yield (**Table 1**). It should also be noted that it is not just ATP turnover that is highest during methylotrophic growth. Strikingly, total Fd_red_ and NADH flux are also about two-fold higher under methanol grown conditions than during cofeeding (**Fig. 6a, b**). We speculate that the need for higher total co-factor turnover flux during growth on methanol may result in a lower proteome efficiency, underlying the increased maintenance costs under these conditions.

It should be pointed out that the cofactor for one step in the WL pathway in *E. limosum* – the methylene tetrahydrofolate reductase (MTHFR) – is not known. The analysis above is based on the assumption commonly used in the field that it is a bifurcating enzyme. Another prominent possibility is that MTHFR forms a complex with Rnf, allowing the reduction of methylene-THF to methyl-THF to be coupled in an energy-conserving manner with oxidation of ferredoxin (Öppinger et al., 2022). We therefore repeated the cofactor analysis with this alternative model, using the carbon fluxes from the original analysis (**Table S14**). Generally, the results support the conclusions above (**Fig. S5**), with glycolysis supplying only a small fraction of the ATP during cofeeding, a 2-fold higher maintenance requirement during methylotrophic growth, and overall higher cofactor turnover rates.

The significant reduction in specific methanol uptake rate compared to cofeeding prompted us to wonder whether this was a result of transcriptional regulation. We thus carried out differential gene expression analysis through RNAseq on samples obtained from each growth condition during chemostat growth (**Tables S15, S16, S17**). As expected based on previous analysis of a methanol-dependent regulator in *E. limosum* (Bae et al., 2026), genes involved in methanol metabolism were highly upregulated on methanol compared to glucose, with the corrinoid-dependent methyltransferase genes showing the highest log_2_-fold changes between these conditions. Interestingly, however, there was no statistically significant difference in expression of these genes between growth on methanol and during cofeeding. This indicates that the decreased methanol uptake during cofeeding results from a post-transcriptional regulatory mechanism. Translational regulation of WLP proteins during autotrophic growth has been previously reported in *E. limosum* (Song et al., 2018).

The above analysis suggests that, by reducing specific methanol uptake, glucose cofeeding avoids ATP- and Fd_red_-intensive steps associated with methanol metabolism, resulting in a lower turnover of all energy cofactors. An unresolved question, however, is which of these resources is most limiting. Ultimately, the largest sink for ATP during methylotrophic growth is the reversal of the Rnf/ATPase to ‘upgrade’ electrons from NADH to Fd_red_. Perhaps, then, the actual limitation is the production of Fd_red_, not ATP specifically. ^13^C-MFA cannot resolve this question, but now that the genetic toolbox for *E. limosum* is well developed, mutational studies could provide interesting insights. For example, replacing the NAD-dependent, ATP-producing GAPDH (*gapA*) with the non-phosphorylating *gapN* from *Streptococcus mutans* (Mallinson et al., 2023) or the Fd-dependent GAPOR from *Methanococcus maripaludis* (Park et al., 2007) would allow manipulation of the ratio of ATP to Fd_red_ produced in glycolysis. Additionally, replacement of the putatively electron bifurcating methylene-THF reductase (which requires Fd_red_ for oxidation of methylene-THF) with the characterized NAD-dependent variant from *Acetobacterium woodii,* would decrease the Fd-dependence of methanol oxidation through the WLP, as was investigated recently in *Clostridium autoethanogenum* (Mendelson et al., 2025). More broadly, genetical manipulation of the cofactors used in key steps in acetogenesis represents a promising, largely untapped approach for improving product yield and further enhancing the benefits of substrate cofeeding (Gasparrini et al., 2025).

## 4. Conclusions

This ^13^C MFA conducted for *E. limosum* expands our quantitative understanding of central carbon metabolism across multiple growth substrates. Key to obtaining statistically acceptable flux measurements here was the use of continuous culture to ensure metabolic and isotopic steady state. While continuous culture has fallen out of favor in recent years due to its technical complexity (especially for anaerobes), advances in low-cost, small-scale reactor systems like AneVO (Hoyt et al., 2026; Wong et al., 2018) allow for these experiments to be conducted with minimal complexity on the benchtop in small-volume cultures ideally suited for work with expensive isotopic substrates. Our analysis revealed two surprising findings: First, that cysteine uptake contributes substantially to central carbon metabolism, resulting in net oxidative flux through PFOR even on substrates thought to require reductive flux for synthesis of gluconeogenic intermediates. Second, although the benefit of sugar cofeeding in other work results from the provision of additional ATP, during methanol/glucose co-metabolism in *E. limosum,* only minimal ATP is derived from glucose catabolism. Instead, the improved growth here stems from a reduced methanol flux through the resource-intensive WLP, resulting in a reduction in the total rate of energy cofactor turnover. Ongoing studies leveraging the advanced genetic toolbox for *E. limosum* will be required to determine which resource – ATP or Fd_red_ – is the major limitation during methylotrophic growth.

Finally, ^13^C-MFA revealed that PFOR operates very close to equilibrium, consistent with the view that acetogens live close to the ‘thermodynamic limit of life’ (Schuchmann and Müller, 2014). We also showed qualitatively that the WLP is highly reversible during growth on methanol. Though we could not precisely quantify this here, future studies that use recently validated techniques to rigorously measure the extent of ^13^C-labeling of key tetrahydrofolate intermediates (Chen et al., 2017) could provide quantitative assessment of exchange fluxes through these reactions. Overall, the quantitative insights from this work will be indispensable for ongoing engineering of *E. limosum* and other acetogens as microbial biocatalysts for C1 conversion.

## Author Contributions

KOH: Conceptualization, Investigation, Methodology, Roles/Writing -original draft

WG: Investigation, Methodology

JAH: Methodology

BMW: Conceptualization, Funding acquisition, Writing - review & editing, Supervision Methodology.

## Acknowledgements

This work was supported by Startup Funds from Northeastern University, the DOE ARPA-E ECOSynBio program (Award #DE-AR0001511), the National Science Foundation (Awards #2345872, #2441203). KOH was additionally supported by the Ford Foundation Predoctoral Fellowship.

## Supporting Information

Supporting information can be found XXXXXX

