## Supplementary Information for "^13^C-Metabolic Flux Analysis of Substrate Co-Metabolism in the Methylotrophic Acetogen *Eubacterium limosum*"

**Figure S1. Benchtop Anaerobic Chemostat Setup**

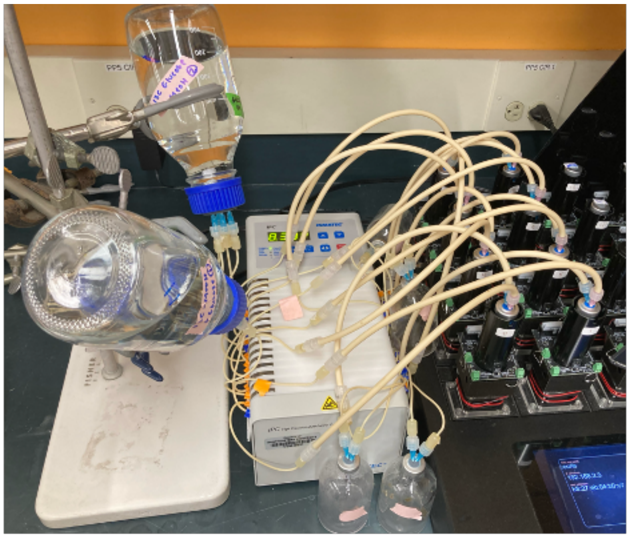

**Table S1. Growth data for cultures with alternative reducing agents**

| **Reducing Agent** | **Lag Time (h)** | **Final OD (N=1)** |
|---|---|---|
| 2 mM $Na_2S$ | 300 h | 0.3 |
| 1 mM $Na_2S$ | 100 h | 0.2 |
| 0.5 mM $Na_2S$ | 100 h | 0.3 |
| 2 mM $Na_2S$ + 1 mM Met | 300 h | 0.2 |
| 1 mM $Na_2S$ + 1 mM Met | 100 h | 0.2 |
| 0.5 mM $Na_2S$ + 1 mM Met | 200 h | 0.2 |
| 3 mM Cysteine | 50 h | 1.5 |

**Figure S2. Cysteine is consumed during growth on either methanol or glucose.**

Mass Isotopologue Distributions (MID) for proteinogenic amino acids obtained from *E. limosum* grown with [3-$^{13}$C] cysteine, with methanol (a.) or glucose (b.). Standard deviation for MID measurements were less than 1% for glucose grown, and less than 6% for methanol grown.

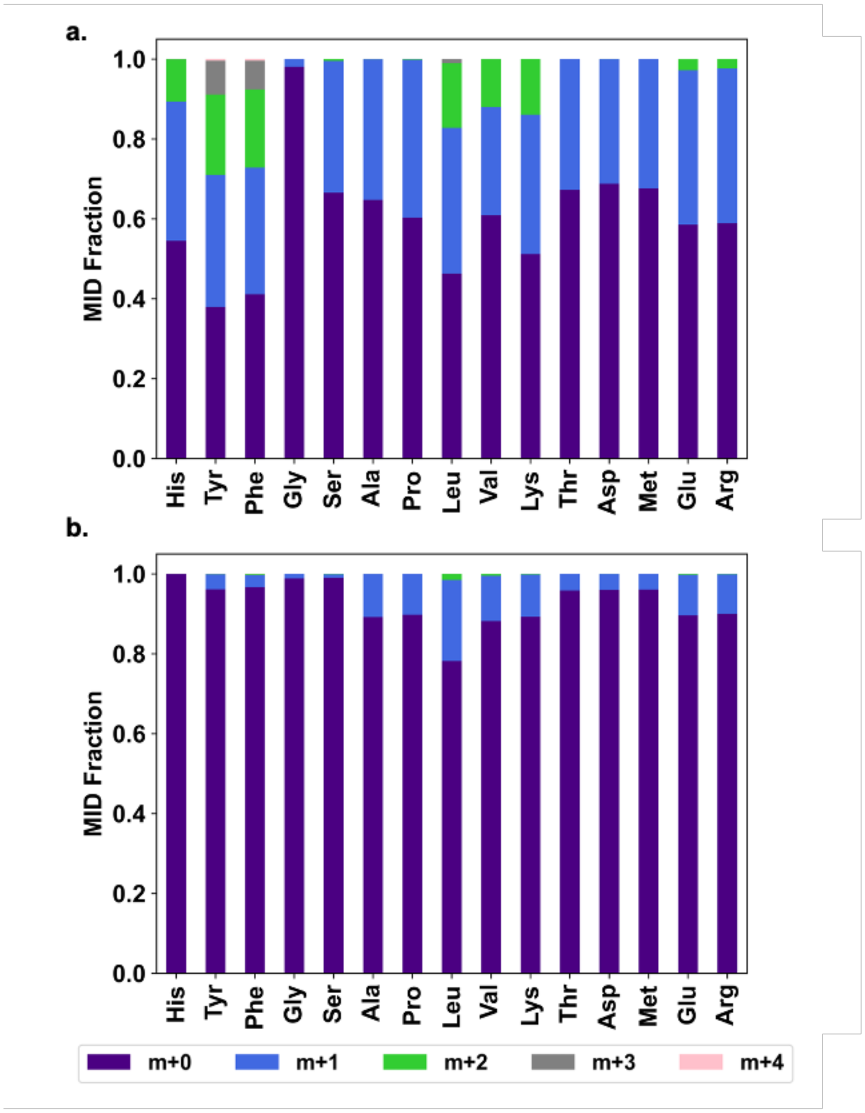

**Figure S3. Methanol-Glucose Cofeeding Pilot tests in AneVO chemostat.**

Each chemostat was initiated with a 0.02 $h^{-1}$ dilution rate at 330 hours, then increased to 0.04 $h^{-1}$ at 568 hours, then increased to 0.08 $h^{-1}$ at 641 hours for the single replicate.

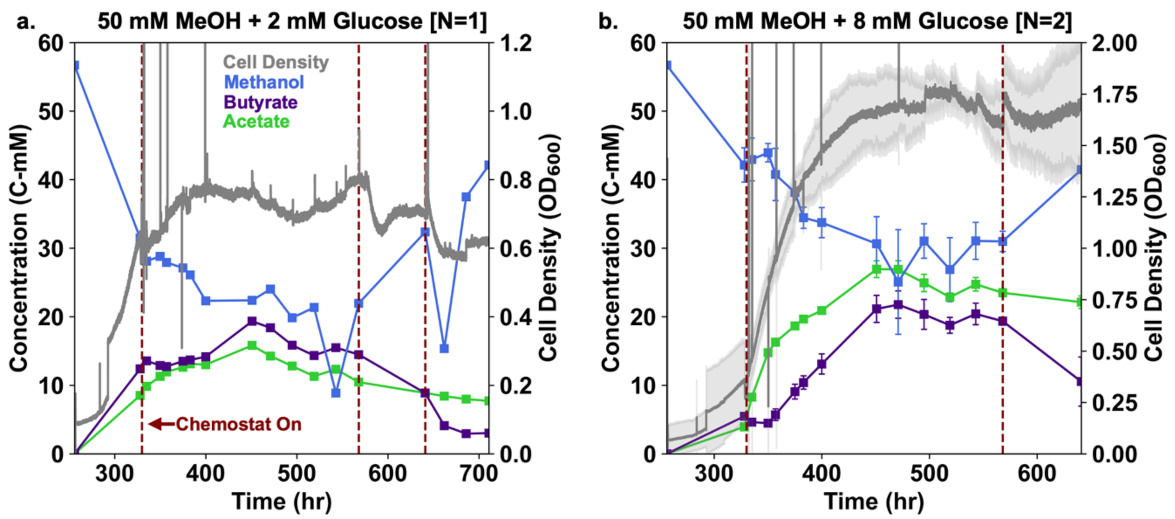

**Figure S4. Data plots for individual culture vials grown on methanol in the AneVO chemostat.**

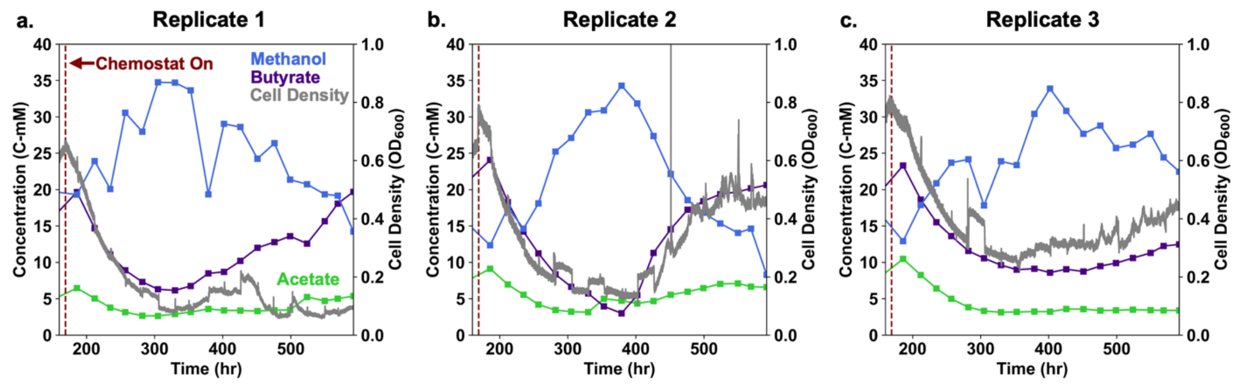

**Table S5. Results of Chi Square Test of MFA**

| Substrate Condition | Number of independent measurements | Number of free fluxes in model | Degrees of Freedom | Upper Tail Chi Square Critical Value for $X_{0.05}$ | Residuals Calculated by Openflux |
|---|---|---|---|---|---|
| Methanol/ $CO_2$ | 321 | 37 | 284 | 324 | 163 |
| Cofed | 324 | 38 | 286 | 326 | 111 |
| Glucose/ $CO_2$ | 420 | 36 | 384 | 431 | 272 |

Threshold value is upper tail chi square critical value. Results represent 500 model run iterations in Openflux. Each substrate condition was conducted in triplicate chemostat runs, except for glucose/$CO_2$ which was in quadruplicate.

**Figure S5. Cofactor balances for alternative scenario of non-bifurcating MTHFR.**

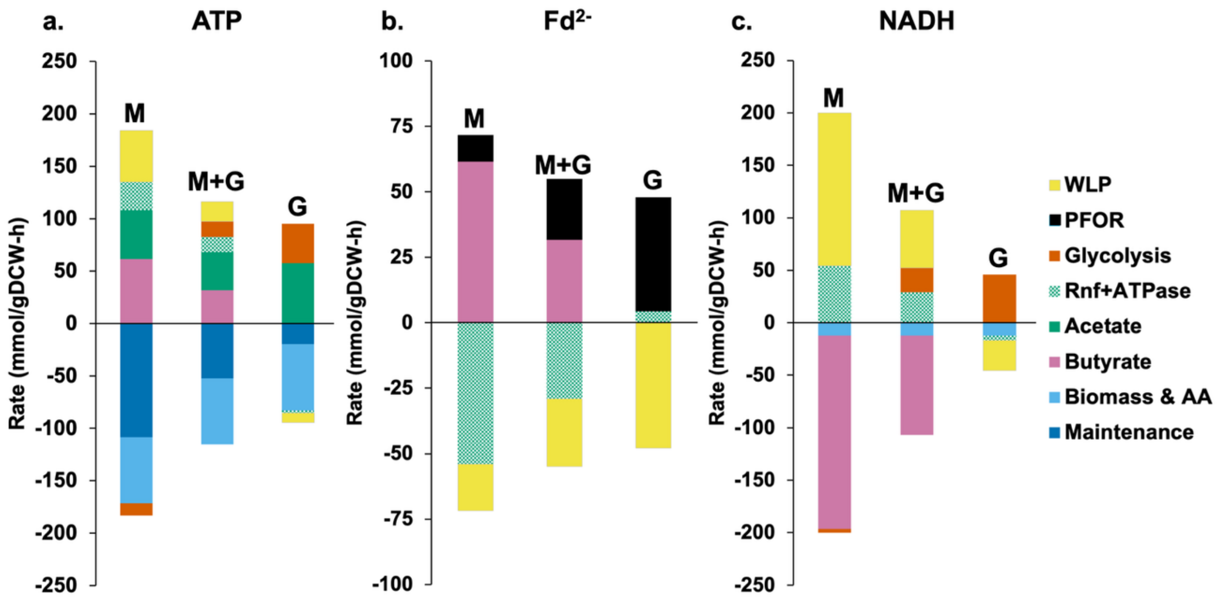
